# Tree diversity and mycorrhizal associations jointly regulate seasonal canopy cooling in forests

**DOI:** 10.64898/2026.07.29.741497

**Authors:** Soroor Rahmanian, Claudia Guimarães-Steinicke, Yuanyuan Huang, Clemens Mehlhorn, Julius Quosh, Olga Ferlian, Hannes Feilhauer, Nico Eisenhauer

## Abstract

The increasing frequency and intensity of heatwaves under climate change highlight the need to understand how biodiversity regulates forest canopy thermal dynamics. Although tree diversity can buffer the microclimate, its effects on canopy temperature and the role of mycorrhizal symbioses remain unclear. We addressed this question in the MyDiv tree diversity experiment in Germany, where tree species richness (1-, 2-, and 4-species mixtures) and mycorrhizal types (arbuscular, ectomycorrhizal, and mixed) are factorially manipulated. During the 2024 growing season, we conducted nine uncrewed aerial vehicle (UAV) surveys using integrated thermal and LiDAR sensors to quantify canopy temperature and structural complexity, together with measurements of leaf water content, specific leaf area, soil moisture, and vapour pressure deficit (VPD). Increasing tree diversity generally reduced canopy temperature, although the strength of this relationship varied seasonally and among mycorrhizal types. Cooling effects were strongest during peak summer heat and were more pronounced in arbuscular mycorrhizal (AM) communities than in ectomycorrhizal (EM) and mixed (AM+EM) communities. In contrast, EM communities exhibited greater canopy structural complexity, whereas AM communities maintained higher soil and leaf water content. Structural complexity increased with tree diversity but did not necessarily result in greater canopy cooling. Structural equation modelling revealed that forest thermal buffering emerged through complementary structural and hydraulic pathways, whose relative importance shifted seasonally, with hydraulic regulation becoming increasingly important under hotter and drier conditions. By linking canopy temperature, canopy structure, and plant water relations, our study provides mechanistic insights for understanding how multiple facets of biodiversity regulate forest thermal buffering under climate warming.

## 1 Introduction

Climate change is increasing the frequency, intensity, and duration of heat extremes, posing growing challenges for forest functioning and resilience (IPCC, 2023; Yuan et al., 2019; Novick et al., 2024). Detecting vegetation stress before visible symptoms of decline emerge remains a major challenge, because physiological responses to heat and drought often precede reductions in growth, canopy damage, and mortality. Canopy temperature has emerged as a powerful integrative indicator of plant stress, because it reflects the balance between absorbed radiation, sensible heat dissipation, and evaporative cooling. Consequently, canopy temperature is closely linked to plant water status, stomatal regulation, photosynthetic performance, and drought responses, providing an early indication of vegetation stress and ecosystem vulnerability (Jones et al., 2014; Urban et al., 2017; Still et al., 2021; Still et al., 2022). Because elevated canopy temperatures can impair carbon uptake, increase hydraulic stress, and accelerate mortality risk, understanding the ecological mechanisms regulating canopy temperature is critical for predicting forest resilience under climate warming.

Biodiversity is increasingly recognized as a key determinant of ecosystem resistance to climatic stress (Isbell et al., 2015; Anderegg et al., 2018; Sasaki et al., 2026). Diverse plant communities can modify microclimatic conditions by altering canopy structure, evapotranspiration, and turbulent exchange, often reducing temperatures and atmospheric demand (Bonan, 2008; De Frenne et al., 2021; Zellweger et al., 2020). Greater structural complexity may enhance canopy– atmosphere coupling and heat dissipation by increasing canopy roughness and vertical heterogeneity (Hardiman et al., 2013; Ehbrecht et al., 2017; Atkins et al., 2018). Consistent with these mechanisms, recent studies suggest that biodiversity-driven cooling occurs across a wide range of ecosystems (Guimarães-Steinicke et al., 2021; Huang et al., 2024; Wright et al., 2026). While the effects of biodiversity on microclimate have been increasingly documented, the mechanisms by which biodiversity influences canopy thermal dynamics remain poorly understood. In particular, the relative roles of canopy structural complexity and hydraulic regulation, and how their importance shifts across seasons, remain unresolved (De Frenne et al., 2021; Maclean et al., 2021).

Belowground symbioses may provide an additional, but largely overlooked, dimension of forest thermal regulation. Mycorrhizal fungi influence nutrient acquisition, plant-water uptake, and hydraulic functioning, thereby shaping transpiration and responses to atmospheric stress (Augé, 2001; Allen, 2007; Bitterlich et al., 2018). Arbuscular mycorrhizal (AM) tree species generally exhibit more acquisitive water-use strategies and sustained transpiration, whereas ectomycorrhizal (EM) tree species tend to exhibit more conservative hydraulic regulation and greater drought resistance (Lehto & Zwiazek, 2011; Ruiz-Lozano et al., 2016; Terrer et al. 2016; Sachsenmaier et al., 2025). These contrasting strategies may arise from differences in how AM and EM fungi influence plant water and nutrient acquisition, root morphology, and stomatal regulation (Chen et al., 2016; Novick et al., 2016; Grossiord et al., 2020). Consequently, stands composed of AM tree species may maintain greater stomatal conductance and transpiration when water is readily available, promoting evaporative cooling through greater partitioning of available energy into latent rather than sensible heat. In contrast, stands composed of EM tree species tend to reduce water loss more rapidly under drought stress, conserving soil moisture and lowering the risk of hydraulic failure, but potentially reducing evaporative cooling and increasing canopy temperatures (Novick et al., 2016; Grossiord et al., 2020). Consequently, AM and EM stands are expected to differ not only in their water-use dynamics but also in the temporal patterns of canopy temperature regulation across seasons and environmental conditions. However, whether mycorrhizal composition modifies biodiversity-driven canopy cooling through these hydraulic pathways remains unknown.

Beyond their effects on plant water relations, tree communities composed of AM and EM tree species differ in functional composition, because tree species belonging to these mycorrhizal groups exhibit contrasting life-history and functional traits (Phillips et al., 2013; Ferlian et al., 2018). These differences may be reflected in canopy structural complexity through variation in tree height, crown architecture, foliage distribution, and vertical heterogeneity (Ehbrecht et al., 2017; Maeda et al., 2025; Schnabel et al., 2025), although these attributes are also strongly influenced by species identity (Poorter et al., 2006). Consequently, variation in mycorrhizal community composition may coincide with differences in canopy structural complexity, providing a potential structural pathway contributing to forest thermal regulation.

Canopy structure and plant hydraulic functioning are not static but change markedly over the course of the growing season, with important consequences for forest thermal regulation (Polgar & Primack, 2011; Novick et al., 2016). During spring, leaf emergence and canopy expansion progressively increase leaf area and vertical canopy complexity, enhancing light interception while transpiration is generally supported by abundant soil moisture (Polgar & Primack, 2011). As the growing season advances, increasing atmospheric evaporative demand together with declining soil moisture can constrain stomatal conductance and transpiration, thereby reducing evaporative cooling despite a fully developed canopy (Novick et al., 2016; Grossiord et al., 2020). Structural attributes such as canopy height, foliage distribution, and canopy rugosity may therefore become increasingly important for mediating radiation interception, turbulence, and canopy–atmosphere coupling under hot and dry conditions, while hydraulic functioning determines the extent to which trees can sustain transpirational cooling (Hardiman et al., 2013; Ehbrecht et al., 2017; Atkins et al., 2018). These seasonal dynamics differ among tree communities, because species vary in their phenology, canopy architecture, and water-use strategies, and because mycorrhizal types influence plant responses to drought (Polgar & Primack, 2011; Grossiord et al., 2020). Consequently, the relative contributions of structural and hydraulic mechanisms regulating canopy temperature are expected to vary throughout the growing season. Although the structural and hydraulic controls of canopy temperature are increasingly well understood, their relative importance in driving biodiversity effects on canopy thermal regulation remains unclear.

Here, we investigated how tree diversity and mycorrhizal types jointly regulate canopy temperature dynamics in the MyDiv forest biodiversity experiment in Germany, where tree species richness (1-, 2-, and 4-species mixtures) and mycorrhizal types (arbuscular, ectomycorrhizal, and mixed) are factorially manipulated (Ferlian et al., 2018; Weigelt et al., 2021). Using a multi-sensor uncrewed aerial vehicle (UAV) platform equipped with thermal and light detection and ranging (LiDAR) sensors, we quantified canopy temperature and canopy structural complexity across nine campaigns during the 2024 growing season, covering a large gradient in climatic conditions. Structural complexity was characterized by considering canopy height, rugosity, and foliage height diversity, capturing complementary dimensions of canopy vertical organization and heterogeneity. We combined these measurements with assessments of leaf-water content, specific leaf area, soil moisture across multiple depths (5, 15, and 55 cm), and vapour pressure deficit (VPD) to evaluate the structural and hydraulic pathways underlying canopy thermal regulation. We hypothesized that (i) increasing tree diversity reduces canopy temperature most strongly during periods of peak summer heat, (ii) AM-dominated communities exhibit stronger canopy cooling than EM-dominated communities, because acquisitive hydraulic strategies promote greater water availability and evaporative cooling, and (iii) biodiversity-driven canopy cooling emerges through interacting structural and hydraulic pathways whose relative importance shifts across seasons. By integrating canopy thermal dynamics, canopy structure, plant water relations, and mycorrhizal types, our study moves beyond documenting biodiversity-driven cooling to identify the mechanisms through which aboveground and belowground dimensions of biodiversity jointly regulate forest thermal buffering under climate warming.

## 2 Materials and Methods

### 2.1 Study site and experimental design

The study was conducted within the MyDiv biodiversity–ecosystem functioning experiment located at the Bad Lauchstädt Experimental Research Station of the Helmholtz Centre for Environmental Research–UFZ, Saxony-Anhalt, Germany (51°23′ N, 11°53′ E; 114–116 m a.s.l.; Fig. 1). The experiment was established in March 2015 on former arable land characterized by a temperate oceanic climate, with a mean annual temperature of 8.8°C and mean annual precipitation of 484 mm. Soils are fertile Haplic Chernozems developed from loess, with pH ranging from 6.6 to 7.4. Detailed site characteristics are described in Ferlian et al. (2018). The experiment factorially manipulated tree species richness (1-, 2-, and 4-species mixtures) and mycorrhizal composition (AM, EM, and mixed communities (AM+EM)), resulting in eight treatment combinations with ten replicates each.

**Figure 1.**
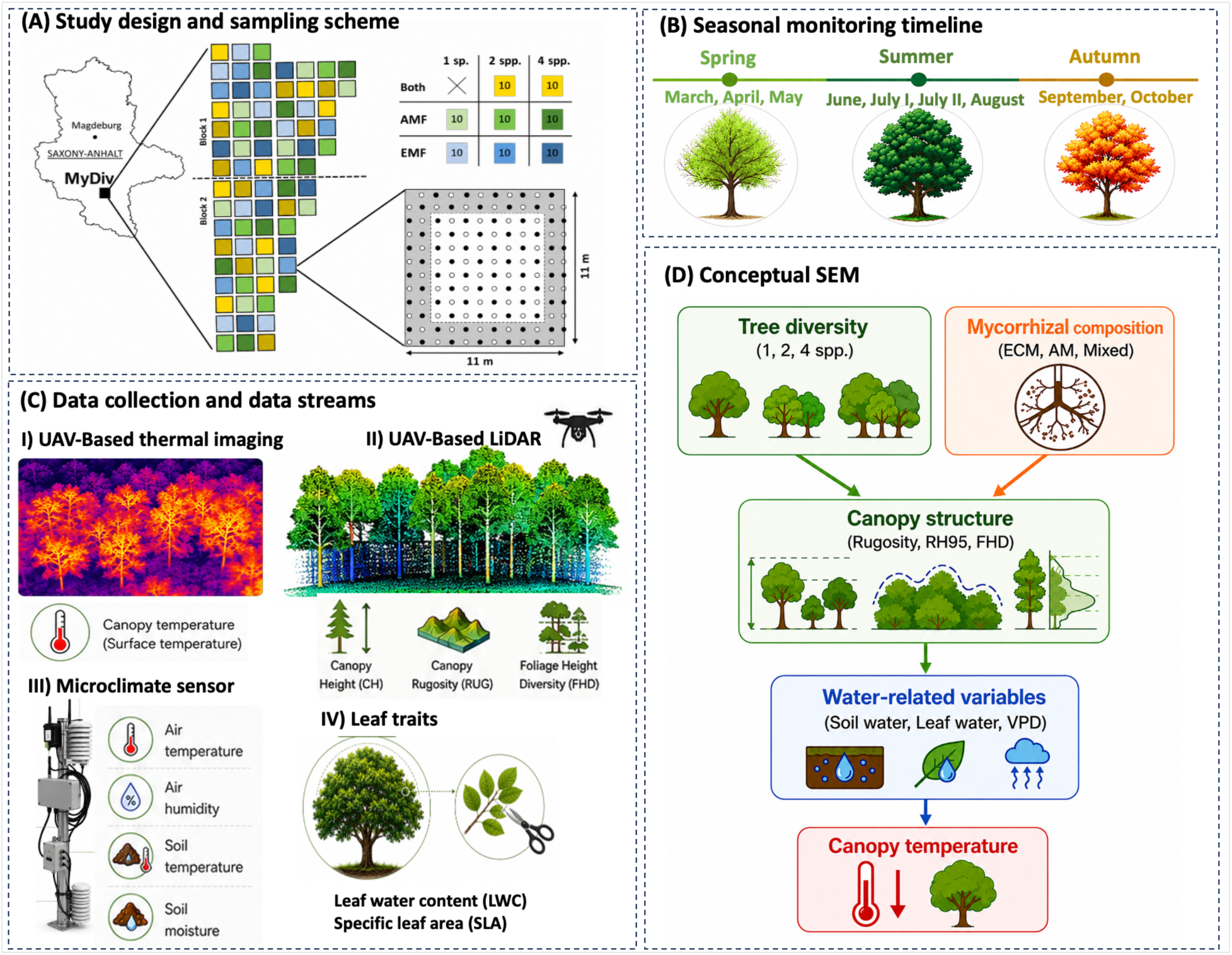
Overview of the study design, seasonal monitoring, data collection, and conceptual framework. **(A)** Study design and sampling scheme at the MyDiv experimental site, including tree species richness treatments (1, 2, and 4 species), mycorrhizal types (AM, EM, and mixed), and plot layout. Panel A was adapted from Ferlian et al. (2018), with graphical modifications. **(B)** Seasonal monitoring timeline covering spring (March–May), summer (June–August), and autumn (September–October). **(C)** Data collection and data streams, including UAV-based thermal imaging for canopy surface temperature, UAV-based LiDAR for canopy structural metrics (canopy height, rugosity, and foliage height diversity), microclimate sensors for air temperature, air humidity, soil temperature, and soil moisture, and leaf trait measurements, including leaf-water content (LWC) and specific leaf area (SLA). **(D)** Conceptual structural equation model (SEM) illustrating the hypothesized associations among tree diversity, mycorrhizal composition, canopy structure, water-related variables (soil water, leaf-water, and vapor pressure deficit [VPD]), and canopy temperature. Some graphical elements in this figure were generated with the assistance of ChatGPT (OpenAI). The figure concept, layout, scientific content, and final composition were developed and verified by the authors.

The experiment comprises 80 plots (11 m × 11 m) arranged in two blocks and planted with ten native temperate deciduous tree species differing in their dominant mycorrhizal types (Figure 1). Five species predominantly associate with arbuscular mycorrhizal (AM) fungi: *Acer pseudoplatanus*, *Aesculus hippocastanum*, *Fraxinus excelsior*, *Prunus avium*, and *Sorbus aucuparia*; and five species predominantly associate with ectomycorrhizal (EM) fungi: *Betula pendula*, *Carpinus betulus*, *Fagus sylvatica*, *Quercus petraea*, and *Tilia platyphyllos*. Mycorrhizal classification was based on published literature (e.g., Wang and Qiu, 2006) and verified through morphological and molecular analyses of roots.

### 2.2 UAV data acquisition

Canopy thermal and structural measurements were collected during nine UAV campaigns conducted between March and October 2024 (19 March, 15 April, 13 May, 10 June, 9 July (July-1), 30 July (July-2), 17 August, 18 September, and 15 October). Surveys were performed under clear-sky and low-wind conditions around solar noon using a DJI Matrice 300 RTK equipped with a Zenmuse H20T thermal sensor and a Zenmuse L1 LiDAR sensor. Thermal imagery was acquired from 35 m above ground level, producing high-resolution canopy temperature maps. LiDAR surveys conducted at the same altitude generated dense three-dimensional point clouds for quantifying canopy structural complexity.

### 2.3 Microclimate measurements

Microclimate sensors installed in the center of each plot, continuously recorded air temperature and relative humidity at 1 m and 15 cm above ground level, soil temperature and volumetric soil water content were measured at depths of 5, 15, and 55 cm at 30-min intervals throughout the study period. Daily mean values were calculated during the UAV campaign date. Vapour pressure deficit (VPD, kPa) was calculated from air temperature and relative humidity following Allen et al. (1998). Saturation vapour pressure was estimated using the Tetens equation (Monteith & Unsworth, 2013), and actual vapour pressure was derived from relative humidity measurements.

### 2.4 Leaf functional traits

Leaf trait measurements were conducted concurrently with UAV surveys during eight sampling campaigns throughout the growing season, with March excluded because canopy development was incomplete. Because processing all plots within the UAV sampling window was not feasible, measurements were restricted to monoculture and four-species mixture plots. For each species within a plot, five to ten fully expanded sun-exposed leaves were collected from different individuals in the upper canopy. Fresh mass was measured immediately after collection. Leaves were subsequently scanned to determine leaf area, saturated in water for 24 h, and oven-dried at 60 °C for 72 h. Specific leaf area (SLA) and leaf-water content (LWC) were calculated following Pérez-Harguindeguy et al. (2013). SLA and LWC were averaged within plots and used to characterize plot-level leaf functional properties.

### 2.5 UAV data processing

#### Thermal imagery

Thermal images were radiometrically calibrated to correct for emissivity, relative humidity, ambient temperature, and sensor-to-target distance effects prior to analysis. The calibrated images were imported into Agisoft Metashape, where images from each sampling campaign were aligned and orthomosaicked into a single georeferenced thermal orthophoto for each time point. To minimize edge effects and thermal contamination from neighbouring plots, analyses were restricted to the central 8 × 8 m core areas within each 11 × 11 m plot. Core areas were delineated in ArcGIS for all 80 plots. To distinguish canopy from non-canopy surfaces, random points were generated within the thermal orthomosaics and assigned to vegetation or soil classes based on visual interpretation. Thermal values extracted from these points were compared using boxplots, and a temperature threshold was determined separately for each sampling date based on the separation of the two distributions. This date-specific threshold was then applied to the thermal orthomosaics to mask soil pixels, including exposed soil and canopy gaps, before extracting plot-level canopy temperature.

#### LiDAR processing

LiDAR data were processed in DJI Terra to generate georeferenced three-dimensional point clouds for each sampling campaign. Canopy structural metrics were subsequently derived from normalized point clouds in R using the *lidR*package (Roussel et al., 2020). Canopy height was characterized by the 95th percentile of vegetation height (RH95), which captures the upper canopy profile and is widely used as a robust indicator of dominant canopy height and forest structural development (Lefsky et al., 2002; Ehbrecht et al., 2017; Maeda et al., 2025). Canopy rugosity was calculated as the standard deviation of canopy heights within each plot, providing an estimate of vertical structural heterogeneity and canopy surface roughness that influences canopy–atmosphere exchange processes (Hardiman et al., 2013; Atkins et al., 2018). Foliage height diversity (FHD) was calculated from the vertical distribution of LiDAR returns using the Shannon diversity index across 1-m height strata. FHD quantifies the diversity and evenness of vegetation occupancy throughout the canopy profile, with higher values indicating greater vertical stratification and structural complexity (Ehbrecht et al., 2017). Together, RH95, rugosity, and FHD capture complementary dimensions of canopy structure, including canopy stature, surface heterogeneity, and vertical complexity.

### 2.7 Statistical analyses

#### Effects of tree diversity, mycorrhizal type, and season on canopy temperature and canopy structure

Linear mixed-effects models (LMMs) were used to evaluate the effects of tree diversity, mycorrhizal type, sampling month, and their interactions on canopy temperature and canopy structural complexity. Tree diversity was included as a log-transformed continuous predictor, whereas mycorrhizal type (AM, EM, and mixed) and sampling month were treated as categorical fixed effects. Separate models were fitted for canopy temperature and each canopy structural attribute (canopy height [RH95], canopy rugosity, and foliage height diversity). Plot identity nested within the block was included as a random effect to account for repeated measurements through time. Temporal autocorrelation among repeated observations was accommodated using a first-order continuous autoregressive correlation structure (corCAR1). Significant interactions were further explored using estimated marginal trends (emtrends) and pairwise comparisons implemented in the emmeans package. Holm-adjusted *P* values were used.

#### Structural equation modelling

Piecewise structural equation modelling (SEM) was used to evaluate the hypothesized pathways linking tree diversity, mycorrhizal type, canopy structure, plant-water relations, VPD, and canopy temperature. Mycorrhizal type was represented as the proportion of AM-associated tree species within each plot (proportion AM), ranging from 0 (EM-dominated) to 1 (AM-dominated). Canopy structural complexity was represented by the first principal component (PC1) derived from canopy height, canopy rugosity, and foliage height diversity. Because these structural metrics were strongly correlated, principal component analysis (PCA) was used to derive a composite structural complexity index; PC1 explained 93.7% of the total variation and was retained for subsequent analyses (Supplementary Fig. S2). Leaf-water content and specific leaf area were included as indicators of plant functional and hydraulic status, whereas shallow soil moisture (5 cm depth) represented belowground water availability and VPD represented atmospheric evaporative demand. Because soil moisture measured at different depths (5, 15, and 55 cm) was highly correlated (Supplementary Fig. S1), only shallow soil moisture was retained in the SEM to avoid multicollinearity. Canopy temperature was specified as the response variable in all models. SEMs were developed based on ecological hypotheses describing how tree diversity and mycorrhizal types influence canopy temperature through interacting structural and hydraulic pathways. Variables that were not significantly associated with other components of the model were removed to improve model parsimony.

Individual pathways were estimated using linear mixed-effects models with plot identity included as a random effect to account for repeated measurements through time. Standardized path coefficients were used to compare the relative strength of relationships among variables. To evaluate seasonal changes in the mechanisms associated with canopy temperature, separate SEMs were fitted for each sampling campaign. August was selected as the representative summer model because it exhibited the strongest biodiversity effects on canopy temperature and the clearest structural and hydraulic relationships associated with canopy cooling. Path coefficients involving leaf-water content were unavailable in March, because canopy foliage had not yet fully developed. Similarly, pathways involving vegetation structure were not estimated in May, because UAV– LiDAR measurements were unavailable owing to technical constraints.

All statistical analyses were conducted in R version 4.5.0 (R Core Team, 2025) using the packages piecewiseSEM (v2.3.0.1), nlme (v3.1-168), lme4 (v1.1-37), lmerTest (v3.1-3), emmeans (v1.11.2-8), and tidyverse (v2.0.0). Model assumptions were assessed by visual inspection of residual diagnostic plots.

## 3 Results and Discussion

Understanding the drivers of canopy temperature is critical for predicting forest responses to climate warming. Here, we tested whether tree diversity regulates canopy temperature, whether this effect depends on mycorrhizal type, and the tree structural, soil, and atmospheric water-related pathways through which these effects emerge.

### 3.1 Seasonal dynamics of canopy temperature and water-related variables

Canopy temperature and associated water-related variables showed pronounced seasonal variation (Fig. S3). Canopy temperature increased from spring to summer, peaking in July, and declined toward autumn. VPD followed a similar pattern, indicating elevated atmospheric demand during summer. In contrast, leaf-water content and soil water content declined from spring to mid-summer and partially recovered in autumn. These opposing trends indicate a seasonal decoupling between atmospheric demand and water availability (Grossiord et al., 2020; Novick et al., 2016). Summer conditions were therefore characterized by high canopy temperature, elevated VPD, and reduced plant and soil water status, providing the environmental context in which diversity and mycorrhizal effects were most strongly expressed (Anderegg et al., 2017; Yuan et al., 2019).

Monoculture species exhibited substantial variation in canopy temperature throughout the growing season, with species-specific mean temperatures differing by nearly 5 °C (Fig. S4). Canopy temperature differed significantly among species (*P* < 0.001), and species-specific thermal responses varied across sampling months (species × month: *P* = 0.008). This pronounced interspecific variation indicates that tree species possess distinct thermal characteristics that may contribute to the biodiversity effects observed in mixed-species communities.

### 3.2 Seasonal variation in canopy temperature across tree diversity and mycorrhizal types

Tree diversity significantly reduced canopy temperature, but the magnitude of this relationship varied throughout the growing season (Fig. 2; Table S1). No consistent relationship was observed during early spring, when canopy development was incomplete, or in late autumn following leaf senescence. In contrast, canopy temperature declined with increasing tree diversity from late spring onwards, with the strongest negative relationships occurring during periods of peak summer heat. These seasonal patterns indicate that biodiversity-driven canopy cooling emerged primarily under conditions of fully developed canopies and high atmospheric demand, when transpirational cooling and canopy–atmosphere interactions are expected to be strongest.

**Figure 2.**
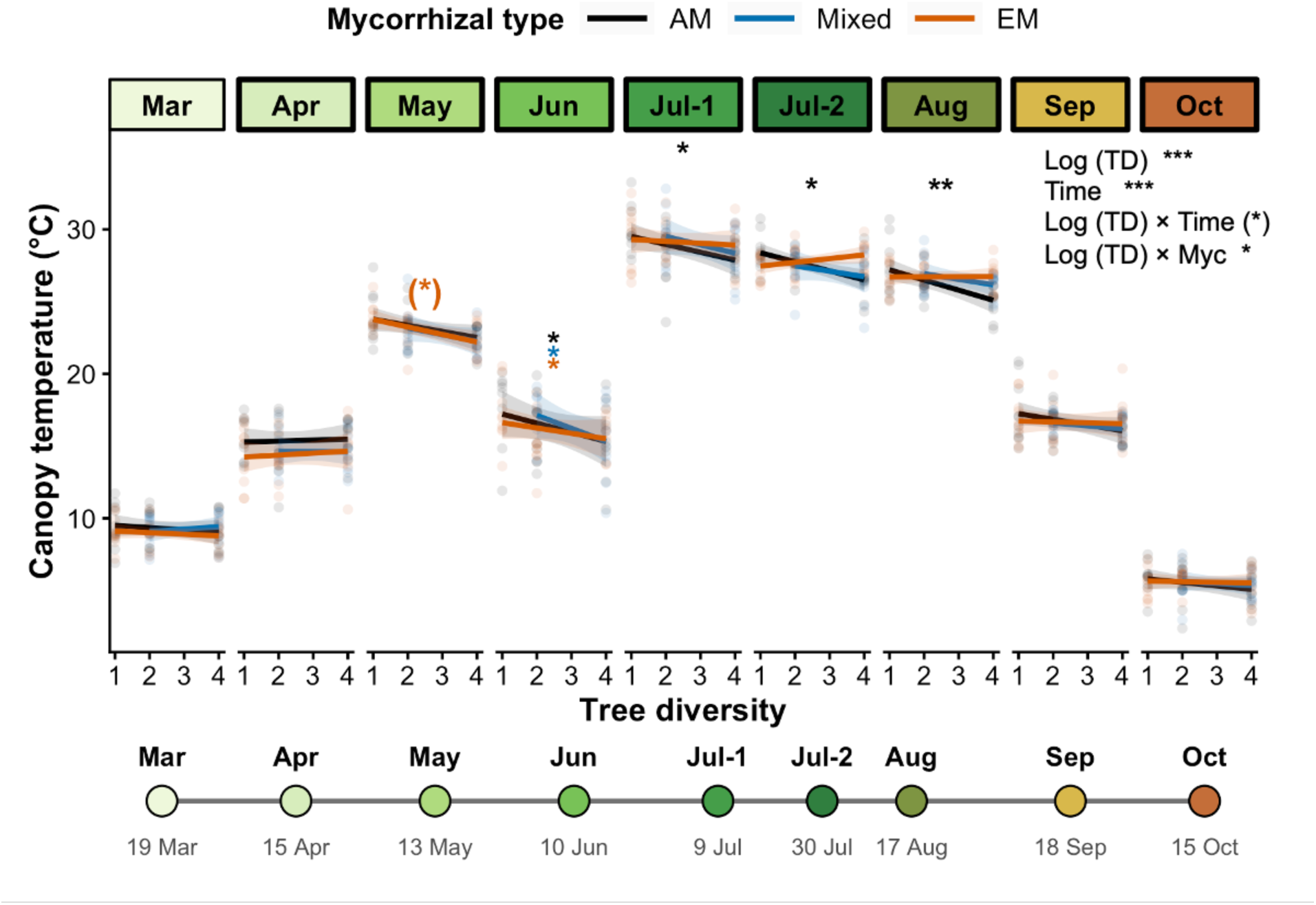
Seasonal relationships between tree diversity, mycorrhizal type, and canopy temperature. Relationships between tree diversity and UAV-derived canopy temperature across nine sampling campaigns from March to October 2024. Points represent individual observations, and coloured regression lines indicate fitted linear relationships for the three mycorrhizal treatments (AM, Mixed, and EM) within each sampling campaign. Tree diversity was log-transformed for the statistical analyses, whereas the x-axis displays the original tree diversity values. Asterisks above the regression lines indicate whether the estimated slope differs significantly from zero within each mycorrhizal treatment (Holm-adjusted *P* values; ns, non-significant; (*), *P* ≤ 0.10; * *P* < 0.05; ** *P* < 0.01; *** *P* < 0.001). The overall significance of the fixed effects from the linear mixed-effects model is summarised in the upper-right corner. Coloured panel headers and the timeline indicate the seasonal progression from spring to autumn. Abbreviations: TD, tree diversity; Myc, mycorrhizal type.

A significant interaction between tree diversity and mycorrhizal type indicated that the relationship between tree diversity and canopy temperature differed among mycorrhizal types (Fig. 2; Table S1). Overall, increasing tree diversity was associated with stronger negative relationships with canopy temperature in AM communities than in EM communities, whereas mixed AM–EM communities exhibited intermediate responses (Fig. 2). These patterns were most apparent during the summer months, when biodiversity-driven cooling was strongest.

These findings indicate that the thermal benefits of increasing tree diversity depended on mycorrhizal type rather than being expressed uniformly across communities. Similar context-dependent effects of mycorrhizal type on biodiversity–ecosystem functioning relationships have previously been reported for forest productivity (Luo et al., 2024).

### 3.3 Effects of tree diversity and mycorrhizal type on seasonal canopy structural complexity

Tree diversity consistently promoted the development of structurally more complex canopies throughout the growing season (Fig. 3). Canopy rugosity, canopy height (RH95), and foliage height diversity (FHD) all increased with tree species richness, indicating that more diverse stands developed taller and vertically more heterogeneous canopies. Positive relationships between tree diversity and canopy structural traits were observed throughout the growing season, with the strongest correlations generally occurring for canopy rugosity and FHD (Fig. 3A, C), whereas canopy height showed weaker but consistently positive relationships with tree diversity (Fig. 3B). These findings demonstrate that increasing tree diversity consistently enhanced canopy structural complexity across seasons.

**Figure 3.**
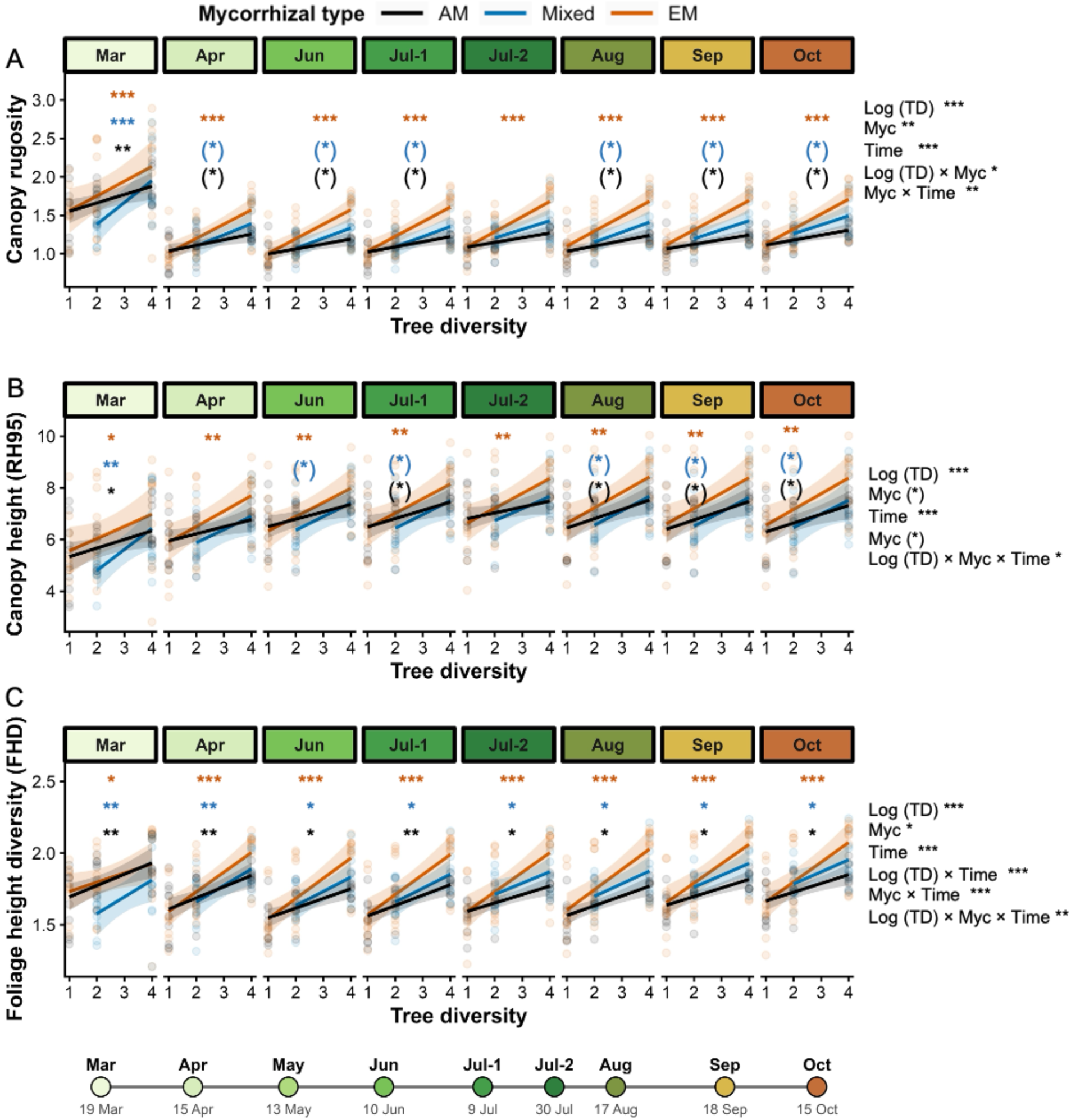
Seasonal relationships between tree diversity and canopy structural complexity across the growing season. Relationships between tree diversity (1, 2, and 4 tree species) in plots dominated by ectomycorrhizal (EM), mixed ectomycorrhizal–arbuscular mycorrhizal (Mixed), and arbuscular mycorrhizal (AM) tree communities and (A) canopy rugosity, (B) canopy height (RH95), and (C) foliage height diversity (FHD) derived from UAV–LiDAR measurements collected during seasonal monitoring campaigns from March to October. No LiDAR-derived canopy structural measurements were available in May owing to technical constraints. Each point represents an individual plot. Solid lines show fitted linear regressions, and shaded bands indicate 95% confidence intervals. coloured asterisks indicate the significance of the simple slope of log-transformed tree diversity within each mycorrhizal type and month, estimated using emtrends with Holm-adjusted P-values. Significance levels are: (*) P < 0.10, * P < 0.05, ** P < 0.01, *** P < 0.001. Abbreviations: TD, tree diversity; Myc, mycorrhizal type

Mycorrhizal type also influenced canopy structural development, although the magnitude of these effects varied among structural traits and throughout the growing season (Fig. 3). Canopy height (RH95) did not differ significantly among mycorrhizal types during any sampling campaign (Fig. 3A). In contrast, canopy rugosity differed among mycorrhizal types from mid-summer onwards, with EM stands generally exhibiting greater rugosity than AM stands, while mixed communities showed intermediate values (Fig. 3B). Differences in foliage height diversity were less consistent and became evident only during late summer and early autumn, when EM stands also tended to exhibit higher FHD than AM stands (Fig. 3C).

The consistent positive effects of tree diversity on canopy structural complexity support the hypothesis that biodiversity enhances the three-dimensional organization of forest canopies (Ray et al., 2023). Increased canopy height, rugosity, and vertical foliage distribution are expected to strengthen canopy–atmosphere coupling, enhance turbulent mixing, and facilitate heat dissipation (Hardiman et al., 2013; Ehbrecht et al., 2017; Atkins et al., 2018). However, structural complexity alone could not fully explain the observed patterns of canopy temperature. Despite exhibiting greater canopy rugosity and, during parts of the growing season, higher foliage height diversity, EM stands did not display stronger canopy cooling than AM stands (Fig. 2). This decoupling between structural complexity and thermal buffering suggests that canopy architecture alone was insufficient to regulate canopy temperature. Instead, structural attributes likely interacted with hydraulic processes, which are examined in the following sections through seasonal changes in soil water availability, leaf-water content, and stomatal conductance.

### 3.4 Seasonal dynamics of leaf and soil water content

LWC and soil water content (SWC) exhibited pronounced seasonal dynamics throughout the growing season (Fig. 4). Across all mycorrhizal types, both variables declined progressively from spring to late summer before partially recovering in autumn, reflecting the seasonal rise and subsequent decline in evaporative demand.

**Figure 4.**
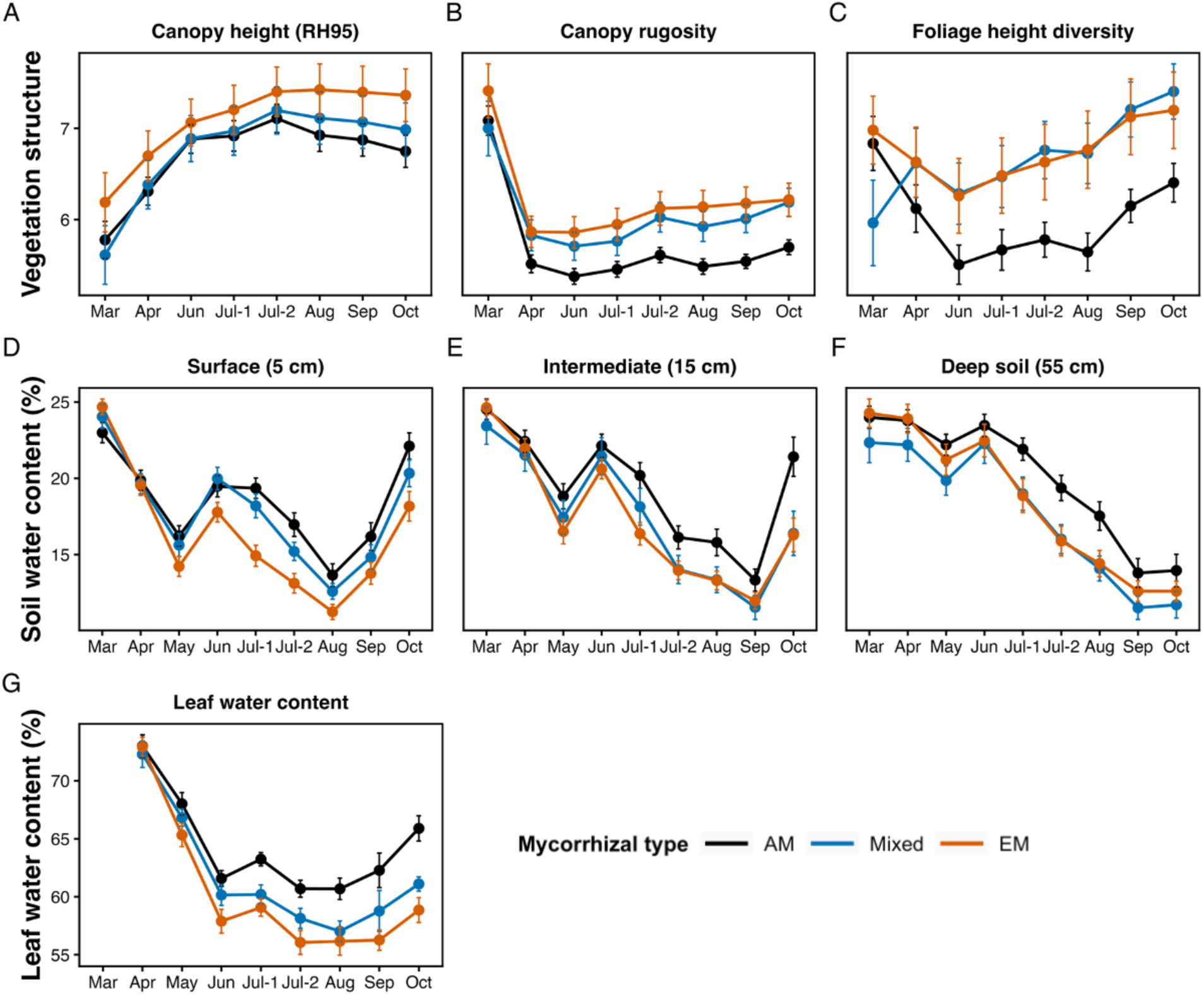
Seasonal variation in vegetation structure and plant–soil water status across mycorrhizal types. (A–C) UAV–LiDAR-derived canopy structural metrics, including canopy height (RH95), canopy rugosity, and foliage height diversity (FHD), measured during eight monitoring campaigns across the 2024 growing season in the MyDiv forest biodiversity experiment. LiDAR-derived canopy structural measurements were unavailable in May owing to technical constraints. (D–F) Soil water content (SWC) at 5, 15, and 55 cm soil depth measured from March to October. SWC values represent the mean measurements collected between 11:00 and 14:00 on each sampling day. (G) Leaf water content (LWC) measured from April to October. Data are shown for plots dominated by arbuscular mycorrhizal (AM), mixed arbuscular–ectomycorrhizal (AM+EM), and ectomycorrhizal (EM) tree species. Points represent plot means ± SE.

Despite this common seasonal pattern, mycorrhizal type consistently influenced ecosystem water status. Leaf-water content remained higher in AM stands than in EM stands throughout most of the growing season, with mixed stands generally exhibiting intermediate values. Differences among mycorrhizal types became most apparent during the summer drought period (July– September), when all communities experienced substantial declines in LWC but AM stands maintained comparatively higher leaf-water status.

Similar patterns were observed for soil water content. Across the three monitored soil depths (5, 15, and 55 cm), soil moisture decreased steadily during summer and reached minimum values between August and September before increasing again in autumn. Although seasonal declines occurred in all treatments, AM stands generally maintained higher soil water contents than EM stands, particularly during the driest part of the growing season, whereas mixed stands again showed intermediate responses.

The close correspondence between seasonal dynamics of soil moisture and leaf-water content indicates strong coupling between belowground water availability and plant water status. These seasonal patterns closely mirrored the stronger biodiversity-driven canopy cooling observed in AM stands during the warmest months (Fig. 2), suggesting that differences in ecosystem water relations contributed to the contrasting thermal responses among mycorrhizal types.

Together with the structural patterns described in the previous section, these results indicate that canopy temperature was jointly regulated by canopy architecture and ecosystem water relations. Although EM stands generally developed greater canopy rugosity and, during parts of the growing season, higher foliage height diversity than AM stands, they did not exhibit stronger canopy cooling (Fig. 4). Instead, the greater leaf and soil water contents maintained by AM stands coincided with lower canopy temperatures despite their comparatively lower structural complexity (Fig. 4). This decoupling between canopy structure and canopy temperature indicates that structural complexity alone was insufficient to explain seasonal variation in canopy thermal regulation.

These findings are consistent with previous studies suggesting that AM fungi may enhance water acquisition and improve root–soil hydraulic connectivity through extensive hyphal networks (Augé, 2001; Phillips et al., 2013). The higher soil and plant water status observed in AM stands may have facilitated greater transpiration and evaporative cooling during the warm season. Conversely, the lower leaf and soil water contents observed in EM stands are consistent with more conservative hydraulic strategies that prioritize water conservation over transpirational cooling (Phillips et al., 2013; Lin et al., 2017; Brzostek et al., 2014). The contrasting structural and hydraulic responses among mycorrhizal types suggest that canopy thermal regulation cannot be explained by canopy architecture alone, but instead emerges from the interaction between structural and hydraulic processes.

### 3.5 Mechanistic pathways of canopy temperature regulation

To explore the mechanisms underlying canopy temperature regulation, we constructed a SEM for August, when differences in canopy temperature among diversity and mycorrhizal types were most pronounced (Fig. 5A). The model explained 38% of the variation in canopy temperature and revealed two interconnected pathways linking biodiversity and mycorrhizal types to canopy thermal dynamics.

**Figure 5.**
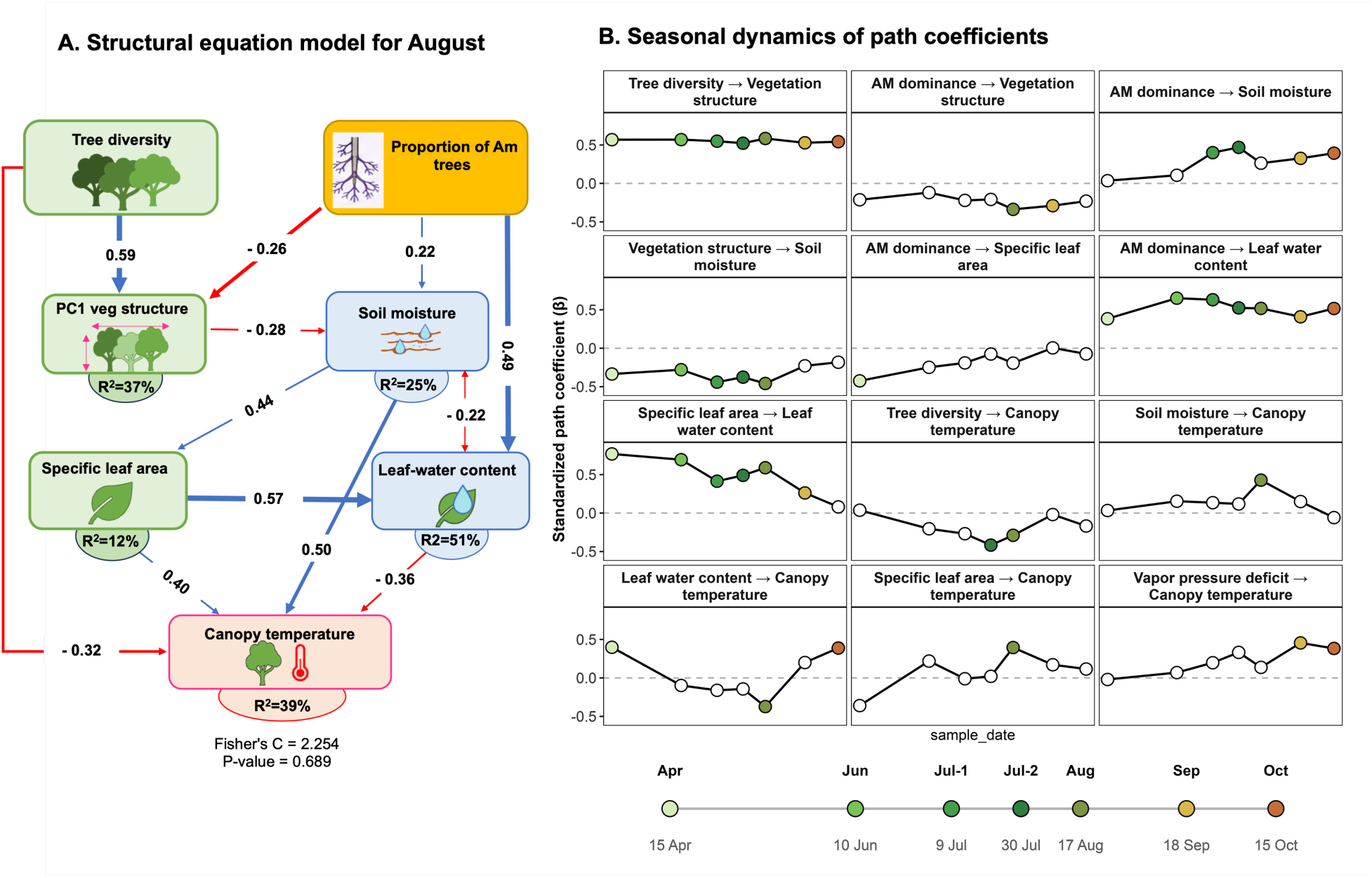
Seasonal structural and hydraulic pathways of canopy temperature regulation. (A) Structural equation model (SEM) for August showing relationships among tree diversity, AM dominance, vegetation structure, specific leaf area (SLA), leaf-water content (LWC), soil moisture, vapour pressure deficit (VPD), and canopy temperature. Vegetation structure represents the first principal component (PC1) derived from canopy height, rugosity, and foliage height diversity. Soil moisture corresponds to shallow mean soil water content (5 cm). Only significant pathways are shown. Blue and red arrows indicate positive and negative standardized path coefficients, respectively, and arrow thickness is proportional to coefficient magnitude. Double-headed arrows indicate correlations. Values within boxes represent explained variance (R²). (B) Seasonal variation in standardized path coefficients for key pathways. Non-significant results are indicated by unfilled symbols. Leaf-water-related pathways were unavailable in March because foliage had not yet fully developed, and vegetation-structure pathways were unavailable in May owing to missing UAV–LiDAR measurements.

The first pathway was associated primarily with tree diversity and vegetation structure. Higher tree diversity was linked to greater structural complexity, which was further associated with variation in specific leaf area and soil moisture. These relationships indicate that biodiversity modifies the physical organization of the canopy and its associated resource environment, forming an indirect structural pathway linking tree diversity to canopy temperature through changes in soil moisture and associated leaf traits. Tree diversity also retained a direct negative association with canopy temperature, suggesting that additional mechanisms not captured by the measured structural and hydraulic variables also contribute to biodiversity-driven cooling.

The second pathway was associated primarily with mycorrhizal type and plant water status. Increasing dominance of arbuscular mycorrhizal (AM) communities was linked to higher soil moisture and greater leaf-water content, whereas increasing ectomycorrhizal (EM) dominance was associated with greater vegetation structural complexity. These contrasting associations suggest that EM-dominated communities were characterized by greater structural complexity, while AM-dominated communities were characterized by improved hydraulic status. In addition, soil moisture and leaf-water content exhibited a significant residual negative covariance, indicating shared variation beyond the directional pathways represented in the model.

The strongest associations with canopy temperature involved leaf-level physiological traits. Higher leaf-water content was associated with lower canopy temperatures, whereas greater specific leaf area was associated with warmer canopies. The positive association between SLA and canopy temperature may reflect the thinner leaves and lower leaf mass per unit area characteristic of high-SLA species (Wright et al., 2004; Poorter et al., 2009), which are associated with differences in leaf energy balance and temperature (Bonan, 2019). Together, these relationships indicate that canopy temperature was closely linked to vegetation water status and leaf functional traits. Notably, the association between AM dominance and higher leaf-water content corresponded with the stronger canopy cooling observed in AM-dominated communities despite their lower structural complexity, suggesting that differences in plant water status may represent an important pathway contributing to thermal buffering during summer.

Although soil moisture showed a positive conditional association with canopy temperature in the August SEM, this relationship should not be interpreted as a direct warming effect of soil water. Instead, it likely reflects the conditional nature of the SEM and covariance among environmental drivers after accounting for the stronger association between leaf water status and canopy temperature (Grace, 2006; Shipley, 2016).

Seasonal analyses of standardized path coefficients further showed that several of these relationships persisted throughout the growing season (Fig. 5B). The positive associations of AM dominance with soil moisture and leaf-water content remained relatively consistent across sampling campaigns, as did the positive relationship between tree diversity and vegetation structure. In contrast, pathways directly connected to canopy temperature exhibited stronger seasonal variation. Associations involving leaf-water content were strongest during the warmest months, coinciding with the period when biodiversity-driven canopy cooling was most pronounced. Towards the end of the growing season, the influence of leaf-water content weakened, whereas the positive relationship between VPD and canopy temperature became significant despite declining atmospheric VPD (Fig. S3). This shift suggests that the strongest canopy cooling occurred when well-developed canopy structure coincided with favorable hydraulic conditions, whereas later in the season canopy temperature became increasingly constrained by atmospheric demand rather than water availability alone.

Our findings suggest that forest thermal regulation cannot be understood solely through canopy structural complexity. Although increasing tree diversity consistently enhanced canopy structural complexity, the strongest canopy cooling occurred only when structural development coincided with favourable hydraulic conditions. Together, these results indicate that biodiversity influences canopy temperature through complementary structural and hydraulic pathways (Ray et al., 2023; Schnabel et al., 2025), whose relative importance shifted seasonally in our study.

Notably, the greater structural complexity observed in EM communities did not translate into lower canopy temperatures. Despite exhibiting higher canopy rugosity and foliage height diversity, EM communities remained warmer than AM communities in mixtures during the summer months, whereas AM communities maintained higher soil- and leaf-water content and exhibited stronger canopy cooling. This decoupling between canopy structure and canopy temperature suggests that structural complexity alone was insufficient to maximize thermal buffering. Instead, canopy cooling appeared to depend on the ability of tree communities to maintain favourable plant water status under increasing atmospheric demand, supporting the importance of hydraulic regulation under heat and drought stress (Drake et al., 2018; Sachsenmaier et al., 2025). Species-level analyses further revealed substantial variation in canopy temperature among monoculture species (Fig. S4), highlighting the important role of species identity in shaping canopy thermal properties. Because AM and EM communities differ in both species composition and functional characteristics, the observed differences in canopy structure likely reflect the combined influence of species identity and mycorrhizal community composition rather than direct effects of mycorrhizal symbioses alone (Phillips et al., 2013; Ferlian et al., 2018). Overall, these findings suggest that forest thermal regulation depends not only on tree diversity, but also on the functional composition of belowground symbioses (Yi et al. 2024), their interaction with species identity, and plant hydraulic functioning, indicating that canopy structural complexity alone is insufficient to explain biodiversity-driven thermal buffering.

Although our findings provide a mechanistic framework for understanding biodiversity-driven thermal buffering, several limitations should be acknowledged. First, canopy temperature was derived from UAV-based thermal imagery and therefore represents canopy surface temperature rather than the complete canopy energy balance (Jones, 2018; Still et al., 2021). Second, although our analyses highlight the importance of hydraulic regulation, key physiological processes such as transpiration, sap flow, and canopy energy exchange were not measured directly but inferred from associated variables, including leaf-water content, soil moisture, and atmospheric demand (Monteith & Unsworth, 2013). Finally, our study was conducted in a young forest biodiversity experiment during a single growing season. As forest structure and species interactions continue to develop over time, the relative importance of structural and hydraulic pathways may change as forests mature or experience more extreme climatic conditions. Such approaches will be essential for determining whether the complementary structural and hydraulic mechanisms identified here are broadly applicable across forest ecosystems and future climatic conditions.

## Conclusion

Our findings demonstrate that tree diversity and mycorrhizal type jointly regulate seasonal canopy temperature dynamics through complementary structural and hydraulic pathways. Increasing tree diversity consistently promoted canopy cooling, particularly during periods of peak summer heat, whereas AM communities exhibited stronger cooling than EM communities despite the latter developing more structurally complex canopies. These results demonstrate that greater canopy structural complexity alone was insufficient to maximize thermal buffering and that maintaining favourable plant water status became increasingly important under elevated atmospheric demand. By integrating UAV-based thermal imaging, LiDAR-derived canopy structure, and plant water-related variables across an entire growing season, our study provides a mechanistic framework linking multiple biodiversity facets to forest canopy thermal dynamics. Unlike most previous biodiversity studies, which have focused primarily on below-canopy or soil-surface microclimates, we directly quantified canopy temperature, the thermal environment experienced by leaves where photosynthesis, transpiration, and thermal stress are tightly coupled (Li et al., 2025), and where many important interactions between trees and microbes (Laforest-Lapointe et al., 2017) and herbivores take place (Li et al., 2025). This perspective provides new insights into how biodiversity regulates canopy functioning under increasingly frequent heat extremes. More broadly, our results demonstrate that biodiversity-driven canopy cooling emerges from the interaction between aboveground structural organization and belowground hydraulic functioning rather than from either mechanism acting alone. By identifying how tree diversity, mycorrhizal type, canopy structure, and plant water relations jointly regulate canopy temperature across seasons, this study improves our understanding of the ecological mechanisms underlying forest thermal buffering. These findings highlight the importance of integrating both aboveground and belowground dimensions of biodiversity when predicting forest resilience and informing the design and management of climate-resilient forests under future climate warming.

## Author Contributions

Soroor Rahmanian: Conceptualization, Data collecting, Data analysis, Writing – Review & Editing

Hannes Feilhauer: Conceptualization, Funding acquisition, supervision, Writing – Review & Editing

Claudia Regina Guimarães-Steinicke: Conceptualization; Data analysis, Writing – Review & Editing

Yuanyuan Huang: Conceptualization, Data analysis, Writing – Review & Editing

Julius Quosh: Data collecting, Writing – Review & Editing

Clemens Mehlhorn: Data collecting, Writing – Review & Editing

Olga Ferlian: Coordination, Writing – Review & Editing

Nico Eisenhauer: Conceptualization, Funding acquisition, Coordination, Supervision, Writing – Review & Editing

## Acknowledgements

S.R. acknowledges the research fellowship provided by the Alexander von Humboldt Foundation (no. 346001466). S.R., N.E., and Y.H. acknowledge support by iDiv ([German Research Foundation, DFG]–FZT 118, 202548816) and the DFG (Ei 862/29-1; FOR 5000, 422440326). Field data acquisition for this study was funded by the German Research Foundation (DFG) with grants SCHM2153/2-1 and FE1331/2-1. We thank the UFZ – Helmholtz Centre for Environmental Research for general on-site support. Further support came from the German Centre for Integrative Biodiversity Research (iDiv) Halle-Jena-Leipzig and the Gottfried Wilhelm Leibniz Prize, funded by the Deutsche Forschungsgemeinschaft (DFG, German Research Foundation) – FZT 118, 202548816; Ei862/29-1. The publication of this article was (partially) funded by [DEAL University publication funds]. We used ChatGPT (OpenAI) to refine the language of this manuscript.

## Data Availability Statement

The data and R code generated during this study have been deposited in the MyDiv repository (Dataset ID: 280; <u>MyDiv repository</u>). They are currently available from the corresponding author upon reasonable request and will be made publicly available through the MyDiv repository after the dataset has been published.

