## Supplementary figures 1-4 and table 1 for "Tree diversity and mycorrhizal associations jointly regulate seasonal canopy cooling in forests"

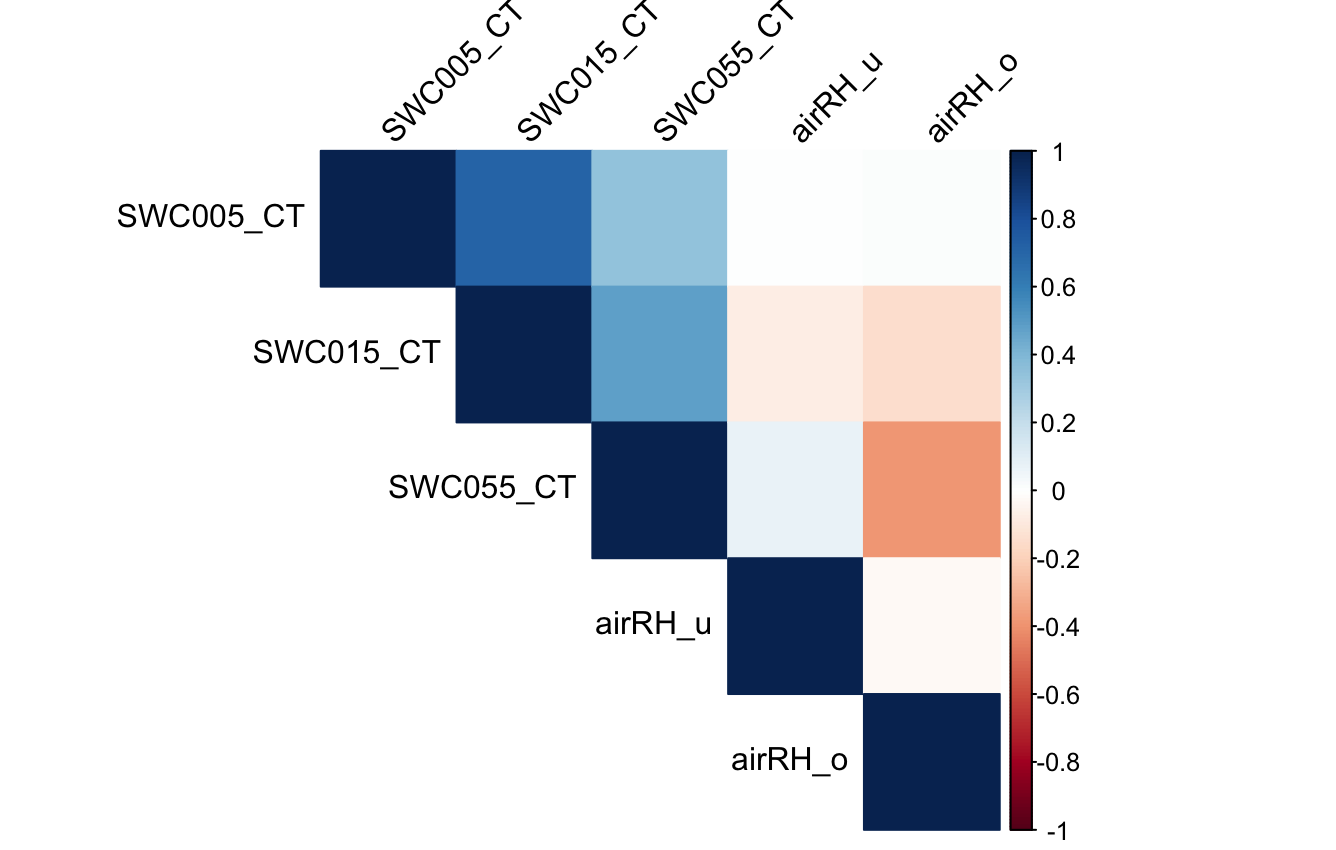

**Figure S1 | Correlation structure of environmental variables.** Pearson correlation matrix showing relationships among volumetric soil moisture measured at 5 cm (SWC005), 15 cm (SWC015), and 55 cm (SWC055) soil depths, and relative humidity measured at 15 cm (airRH_u) and 100 cm (airRH_o) above the ground. Strong positive correlations among soil moisture measurements indicate substantial redundancy across depths. Colors represent correlation coefficients ranging from −1 (red) to +1 (blue).

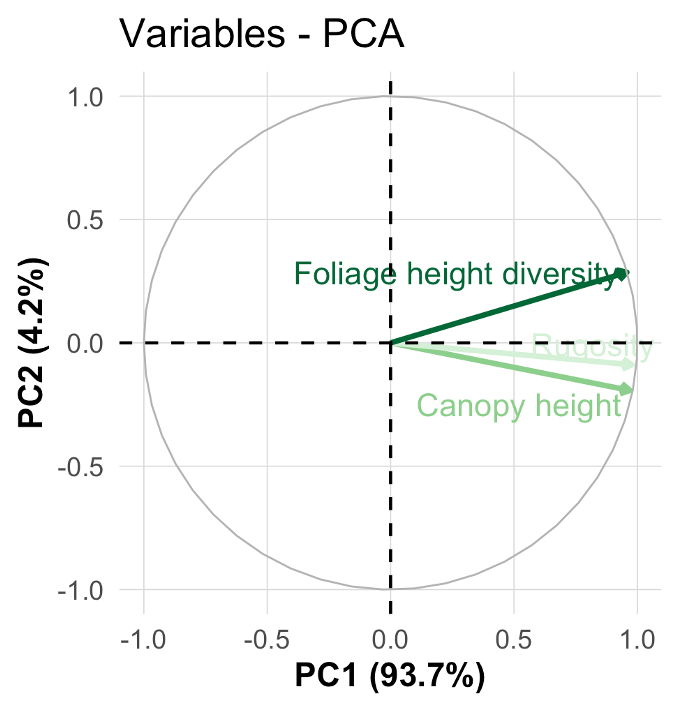

**Figure S2 | PCA-derived canopy structural complexity gradient.** Principal component analysis (PCA) of canopy height (RH95), rugosity, and foliage height diversity (FHD) derived from UAV–LiDAR data. All three metrics loaded strongly and positively on the first principal component (PC1), which explained 93.7% of the total variation and was therefore used as an integrated index of canopy structural complexity in subsequent analyses. The second principal component (PC2) explained 4.2% of the variation. Arrows indicate variable loadings on the ordination axes.

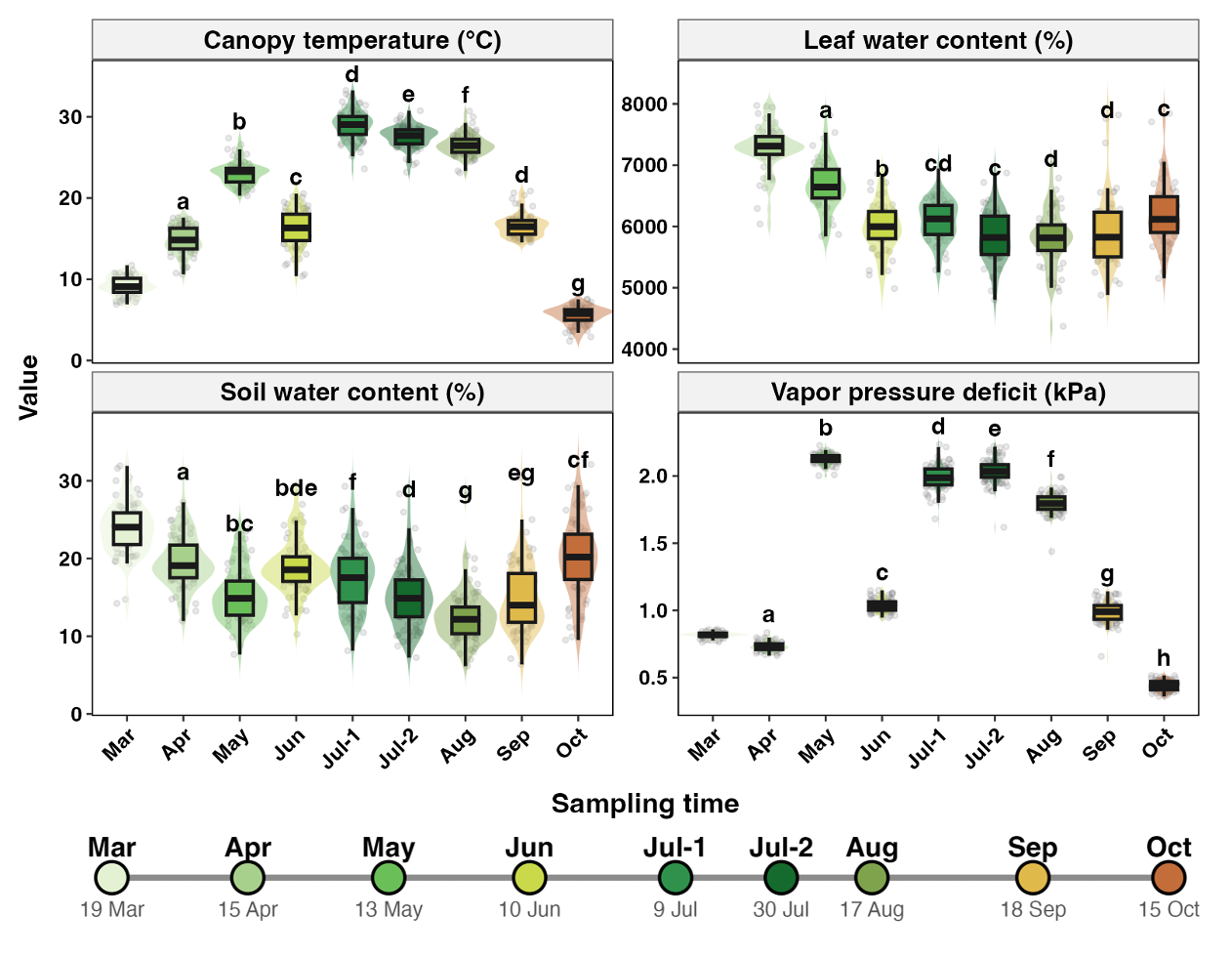

**Figure S3 | Seasonal dynamics of canopy temperature and water-related variables.** Boxplots show variation in canopy temperature, leaf-water content, soil water content, and vapour pressure deficit (VPD) across nine sampling dates from March to October. Points represent individual observations. Letters indicate significant differences among sampling dates based on Benjamini–Hochberg-corrected pairwise Wilcoxon tests. Different colors represent the different sampling dates.

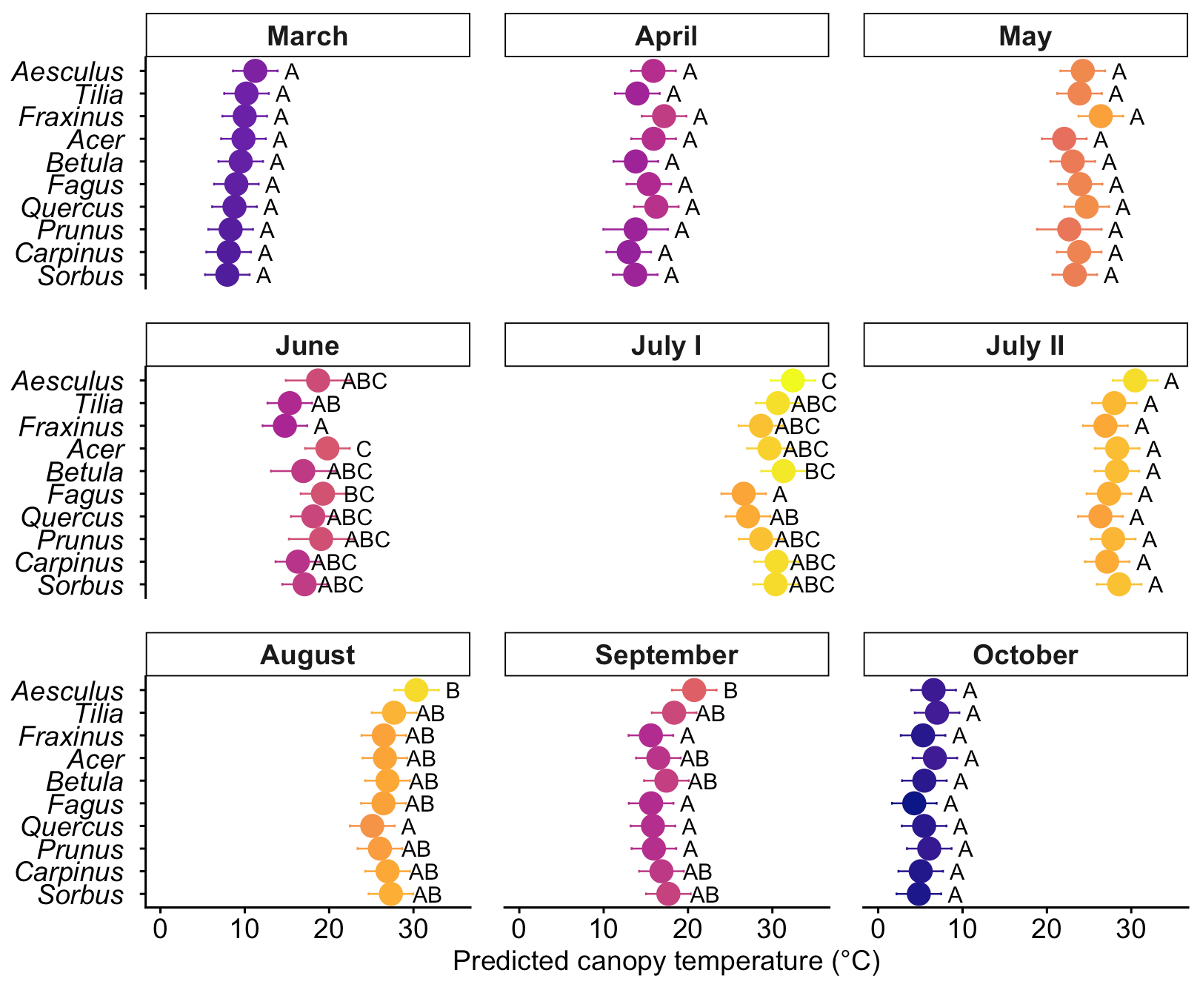

**Figure S4 | Species-specific variation in canopy temperature in monoculture plots across the growing season.** Mean (± SE) canopy temperature measured from UAV-based thermal imagery during nine sampling campaigns (March–October 2024) in monoculture plots of the MyDiv experiment. Points represent species-specific mean canopy temperatures, and error bars indicate standard errors. Different letters indicate significant differences among species within each sampling campaign (Tukey-adjusted pairwise comparisons, *P* < 0.05). Species abbreviations are: *Aesculus hippocastanum* (Aesculus), *Tilia platyphyllos* (Tilia), *Fraxinus excelsior* (Fraxinus), *Acer pseudoplatanus* (Acer), *Betula pendula* (Betula), *Fagus sylvatica* (Fagus), *Quercus petraea* (Quercus), *Prunus avium* (Prunus), *Carpinus betulus* (Carpinus), and *Sorbus aucuparia* (Sorbus). Results demonstrate pronounced seasonal and interspecific variation in canopy temperature under monoculture conditions.

**Table S1.** Linear mixed-effects model testing the effects of log-transformed tree diversity, mycorrhizal type, Time, and their interactions on canopy temperature. The model included nested random intercepts for plots within blocks, specified as (1 | Block/Plot_name).

| Fixed effect | Num. df | Den. df | F | P |
| --- | --- | --- | --- | --- |
| Log (Tree diversity) | 1 | 70.58 | 18.52 | <0.001 |
| Mycorrhizal type | 2 | 70.77 | 2.83 | 0.066 |
| Month | 8 | 574.23 | 474.06 | <0.001 |
| Log (Tree diversity) × Mycorrhizal type | 2 | 70.86 | 3.55 | 0.034 |
| Log (Tree diversity) × Month | 8 | 574.23 | 2.15 | 0.029 |
| Mycorrhizal type × Month | 16 | 574.40 | 0.37 | 0.988 |
| Log (Tree diversity) × Mycorrhizal type × Month | 16 | 574.49 | 0.75 | 0.739 |
